# De novo spatiotemporal modelling of cell-type signatures identifies novel cell populations in the developmental human heart

**DOI:** 10.1101/2021.07.10.451822

**Authors:** Sergio Marco Salas, Xiao Yuan, Christer Sylven, Mats Nilsson, Carolina Wählby, Gabriele Partel

## Abstract

With the emergence of high throughput single cell techniques, the understanding of cellular diversity in biologically complex processes has rapidly increased. The next step towards comprehension of e.g. key organs in the mammal development is to obtain spatiotemporal atlases of the cellular diversity. However, targeted cell typing approaches relying on existing single cell data achieve incomplete and biased maps that could mask the molecular and cellular heterogeneity present in a tissue slide. Here we applied spage2vec, a *de novo* approach to spatially resolve and characterize cellular diversity during human heart development. We obtained well defined spatial maps of tissue samples from 4.5 to 9 post conception weeks, not biased by probabilistic cell typing approaches. We found previously unreported molecular diversity within cardiomyocytes and epicardial cells and identified their characteristic expression signatures by matching them with specific subpopulations found in single cell RNA sequencing datasets.

## INTRODUCTION

Recent efforts to decrypt cellular complexity of human organs, and provide comprehensive maps of their constituent cell types, have been supported by both technological developments and a number of international initiatives^1,2^. One such technological advance is single-cell RNA sequencing (scRNA-seq)^3,4^ enabling profiling the transcriptome of tens of thousands individual cells after tissue dissociation, and thus define the cell-type composition of a tissue architecture. Single cell data can be further combined with more recent spatially resolved techniques^5–9^ to create organ-wide gene expression atlases that map cell-type distributions and spatial biological programs directly in tissue samples, with spatial resolution.

With the aim of producing a spatiotemporal gene expression and cell atlas of the developmental human heart, Asp et al.^10^ recently combined three different high-throughput technologies for gene expression profiling with immunohistochemical staining. They studied three developmental stages in the first trimester at 4.5-5, 6.5 and 9 post-conception weeks (pcw) using (1) Spatial Transcriptomics^9^ (ST) for untargeted spatial gene expression profiling of grid-microdissected tissues, (2) scRNA-seq of 6.5 pcw tissue samples for dissecting cellular heterogeneity at single cell resolution, and (3) in situ sequencing^6^ (ISS) to resolve the spatial heterogeneity at subcellular resolution. Finally, probabilistic cell-mapping via pciSeq^11^ was applied to achieve single cell level maps of the cell-type distribution. PciSeq jointly assigned in situ decoded reads to segmented cells and cells to cell-type gene expression profiles defined with scRNA-seq. The result of the overall study is the first comprehensive spatial atlas of the developmental human heart.

Despite this achievement, some important limitations were found in the study, as pointed out also by Phansalkar et al. ^12^ in a commentary to the paper. One of the main limitations of the probabilistic approach is that preexisting knowledge of the tissue constituent cell-types is required to characterize the spatial cellular heterogeneity. Thus, probabilistic cell typing by in situ sequencing (pciSeq) was only possible for the 6.5 pcw developmental stage where single cell data was available, leaving the cellular diversity in the 4.5-5 and 9 pcw time points unexplored at a single cell level. Additionally, cell-typing methods that depend on priors defined by scRNA-seq, such as pciSeq, may introduce a strong bias that can limit the possibility to distinguish between cell sub-types or sub-states that are not fully resolved by scRNA-seq, as further discussed in the next paragraphs. Finally, most of current cell-typing methods rely on their ability to segment out tridimensional cells from a 2D representation of them, leading to possible misidentification of cells and misclassification of reads.

In order to overcome these limitations and fully explore the spatial heterogeneity of the developmental human heart, we present a *de novo* spatiotemporal analysis of Asp et al. ISS data using all three developmental stages. For our analysis we used a data driven approach, called spage2vec^13^, for generating a spatiotemporal common representation of the spatial gene expression at the different developmental stages. Spage2vec represents the spatial gene expression as a graph and applies a powerful graph representation learning technique to create a lower dimensional representation of the data that is independent from scRNA-seq defined priors. We then used this representation to define the identities and spatiotemporal relationships of cell and sub-cell type gene expression signatures across the three developmental stages of the embryonic heart, identifying previously unreported cell populations within cardiomyocytes as well as in atrial sub-epicardial cells.

## RESULTS

With the aim of exploring cellular diversity during the human heart development, spage2vec^13^ was used to identify cellular expression signatures *de novo* across the three developmental stages **(Figure 1A)** (Methods). The analysis is based on the locations of genes represented in the gene panel used during ISS, and is thus dependent on the panel’s ability to represent heterogeneity in gene expression. A total of 27 clusters with specific cellular expression signatures were found during the heart development **(Figure 1A-B)**. The signatures were grouped in five main classes, according to their expression patterns, including atrial cardiomyocytes, ventricular cardiomyocytes, fibroblast-like cells/epicardium-derived cells, epicardial cells and neural crest cells **(Figure 1B)**. Clusters assigned to the same class were found to have similar molecular and spatial patterns **(Figure 1C-D, Supplementary figure 1-3)**. Their distribution was also found to be consistent through the different samples and time points analyzed and most of them were found to have a conserved location in the heart between pcw 4.5-5, pcw 6.5 and pcw 9 **(Figure 1D, Supplementary figure 1-3)**.

**Figure 1.**
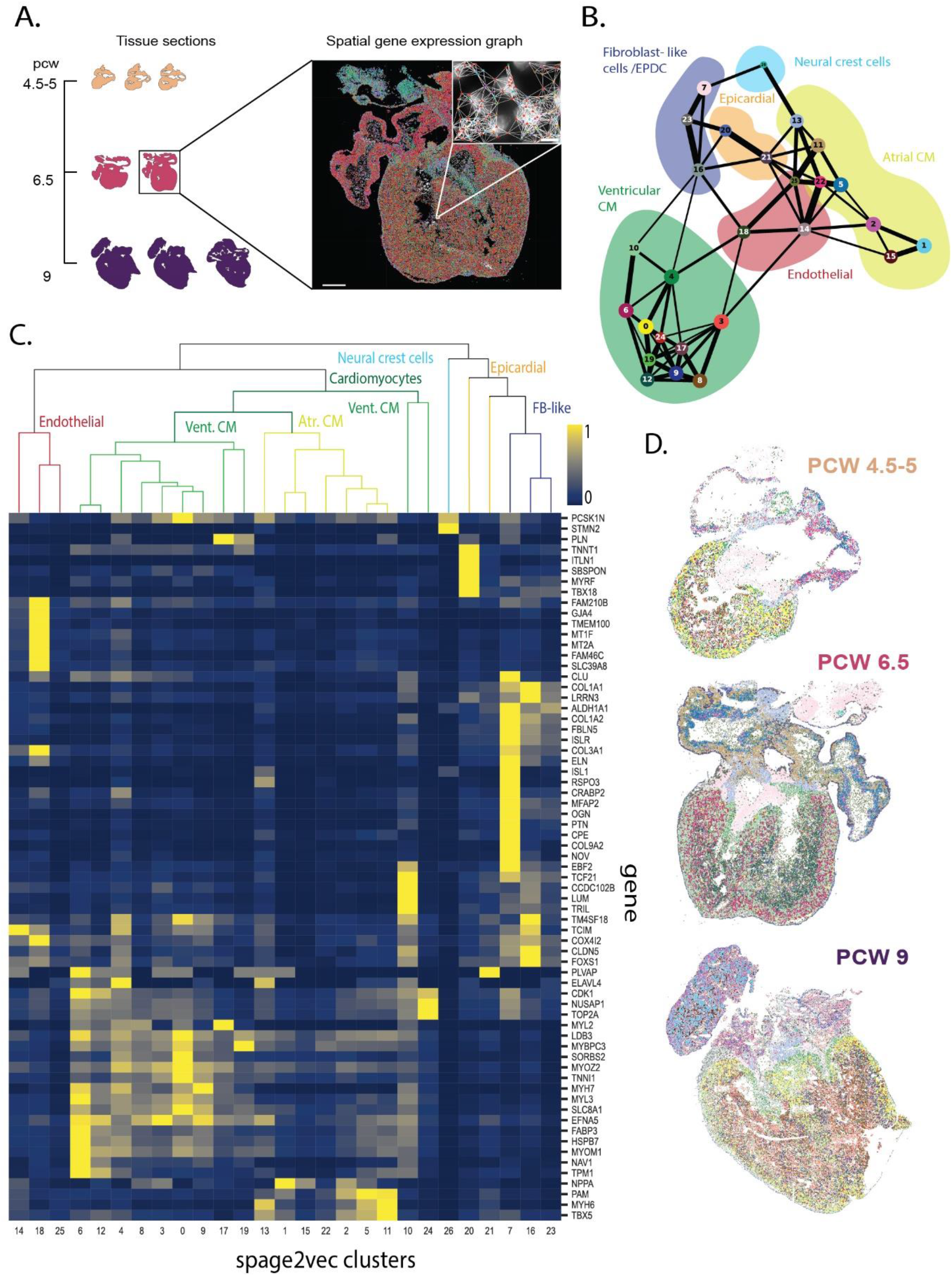
Overview of the Spage2vec approach to characterize the human developmental heart. **A.** Spage2vec constructs a graph from the spatial gene expression of the tissue samples and projects spatial markers in a common latent space. Scale bars: 1 mm, cutout 15 μm. **B.** PAGA plot representing the different expression profiles defined using spage2vec. Background colors represent main cell classes manually annotated based on cluster expression profiles. **C.** Heatmap showing the mean expression of each gene (e.i. expression profile) in the clusters defined by spage2vec, along with suggested cell classes, color coded as in Figure 1C. **D.** Spatial maps of the different expression profiles defined using spage2vec in three sections (color coded as Figure 1B), one from each time point. For interactive multi-resolution viewing: https://tissuumaps.research.it.uu.se/human_heart.html.

In order to identify common findings and discrepancies with previous results, we compared the cellular identities defined by spage2vec with the ones described in *Asp et al*.^10^. This comparison was limited to the only time point analyzed via probabilistic pciSeq^11^; post conception week 6.5 (**Supplementary figure 4**, Methods). A total of 27 meaningful cellular identities shared across all three time points were defined in the spage2vec analysis, in contrast to the 12 cell types defined from scRNA-seq data and assigned in situ via probabilistic cell typing in *Asp et al*. Most of these additional spage2vec clusters capture a previously undescribed diversity within cardiomyocytes **(Supplementary figure 4A)**, while other cell types such as endothelial cells or fibroblast-like cells present a one-to-one correspondence. This is also observed when comparing the expression signatures of the spage2vec clusters and the cell types described in the single cell RNA sequencing dataset from *Asp et al*. **(Supplementary figure 4B)**. Regarding the spatial location of the clusters, both methods agreed on the location of some clusters such as epicardial cells and, to a lesser extent, capillary endothelial cells. However, significant differences were observed when comparing the location of some cell types. This is the case of the clusters with a fibroblast-like expression signature, where spage2vec clusters present a more specific spatial distribution through the tissue in accordance with the previously known location of each of the cell types analyzed and in contrast with the sparser location identified by pciSeq **(Supplementary figure 4C)**.

One of the main concerns of our approach was whether samples with a higher number of cells could be driving the clustering results of the rest of the samples, leading to a misclassification of the cells in the smaller tissue samples. To explore the consistency of the spage2vec clusters found, individual clustering was performed separately on each of the time points (Methods). A total of 94 clusters were found, including 34 in pcw 4.5-5 and 30 both in pcw 6.5 and pcw 9 **(Figure 2)**. Despite small differences, the clusters found in the different time points present a similar distribution in spage2vec latent space for all three time points **(Supplementary figure 5A)**. In addition, most clusters found in specific time points recapitulated molecular and spatial signatures found when analyzing all time points together **(Supplementary figure 5B)**. The clear correspondence between both analyses proves that the diversity found at different time points was not driven by any of the samples individually **(Supplementary figure 6)**.

**Figure 2.**
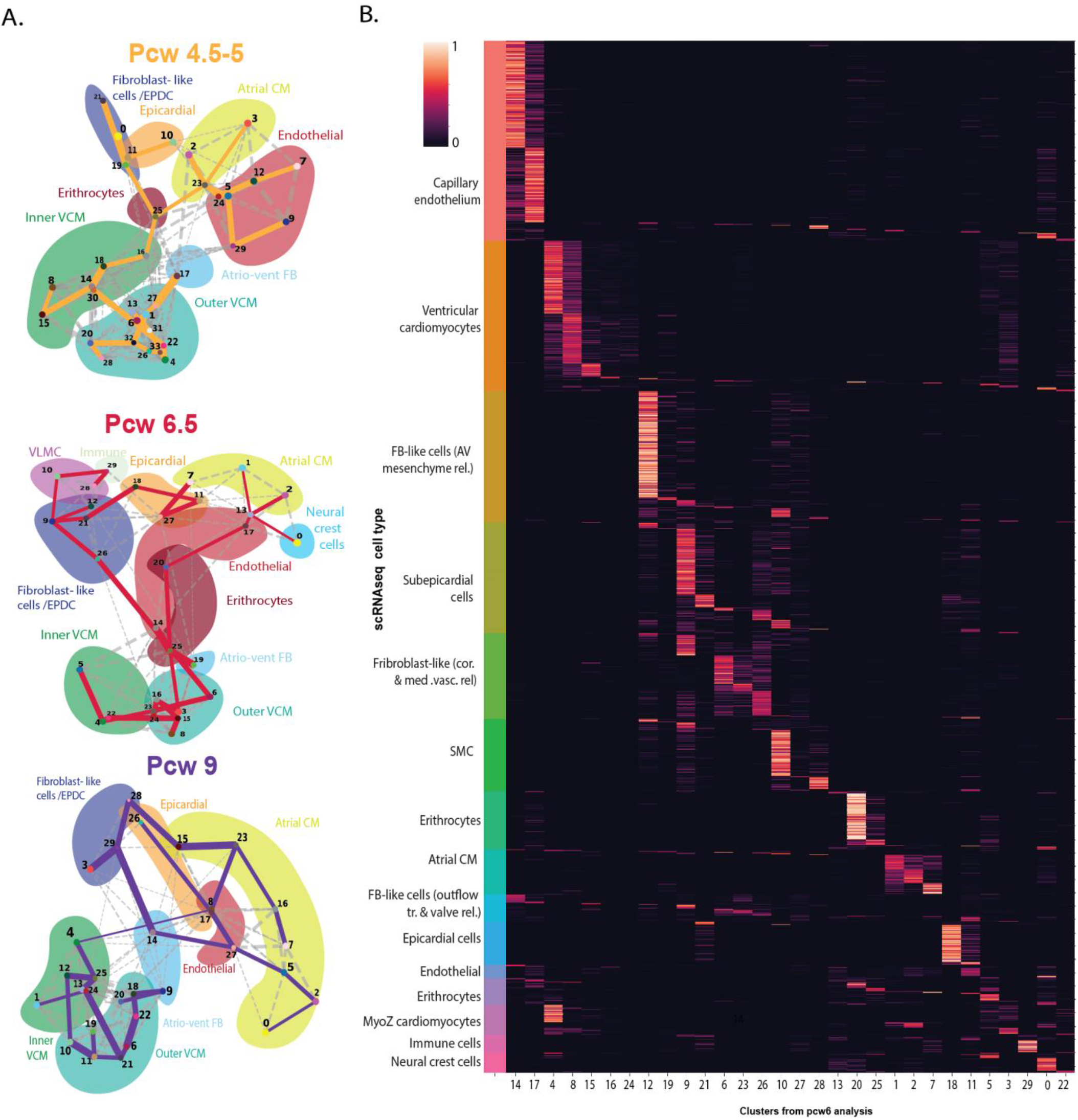
Analysis of individual developmental stages and correspondence with single cell data. A. PAGA plots of the clusters found in each time point specific analysis. Clusters from each time point are represented in a PAGA plot, including 4.5-5 pcw (top), 6.5 pcw (middle) and 9 pcw (bottom). Background colors represent the main cell types found in the dataset. B. Heatmap representing correspondence in terms of cosine similarities between scRNA-seq data and spage2vec clusters from pcw 6.5 (Methods). Cells from scRNA-seq dataset (rows) are sorted based on their cell type in order to facilitate the interpretation.

One surprising aspects of the spage2vec *de novo* analysis is its ability to resolve the cellular heterogeneity at a higher resolution compared to the scRNA-seq data driven analyses, finding a larger number of clusters, that appear to make sense since they show distinct and consistent spatial distribution across the different samples. This may suggest that the spatial organization of biological markers contains essential information for resolving spatial cellular heterogeneity that is masked in scRNA-seq analyses. In order to assess whether traces of this spatially defined diversity could also be found in the scRNA-seq dataset, the molecular signatures from the intermediate time point samples (pcw 6.5) and its corresponding scRNA-seq dataset were integrated using SpaGE^14^ (Method). Correspondence between individual scRNA-seq cells and the different spage2vec clusters defined in pcw 6.5 is shown in **Figure 2B**. Several molecular signatures defined by spage2vec matched specific subpopulations in the single cell dataset, including cardiomyocytes and endocardial cells. With the aim of characterizing these subpopulations, which presented clear spatial locations in the tissue (Supplementary Figure 8), we identified their most differentially expressed genes **(Figure 3A, Supplementary Figure 7)** and assessed their gene ontology (GO) characteristics using scRNA-seq (Methods). With this, we identified two specific endocardial subpopulations: one located in the atria and the other one in the ventricles **(Figure 3B)**. Although they show very distinct spatial localization, we did not find notable differences in the expression of differentially expressed genes, and both show enrichment in GO terms involved in cardiovascular morphogenesis and development **(Fig. 3A, C)**.

**Figure 3:**
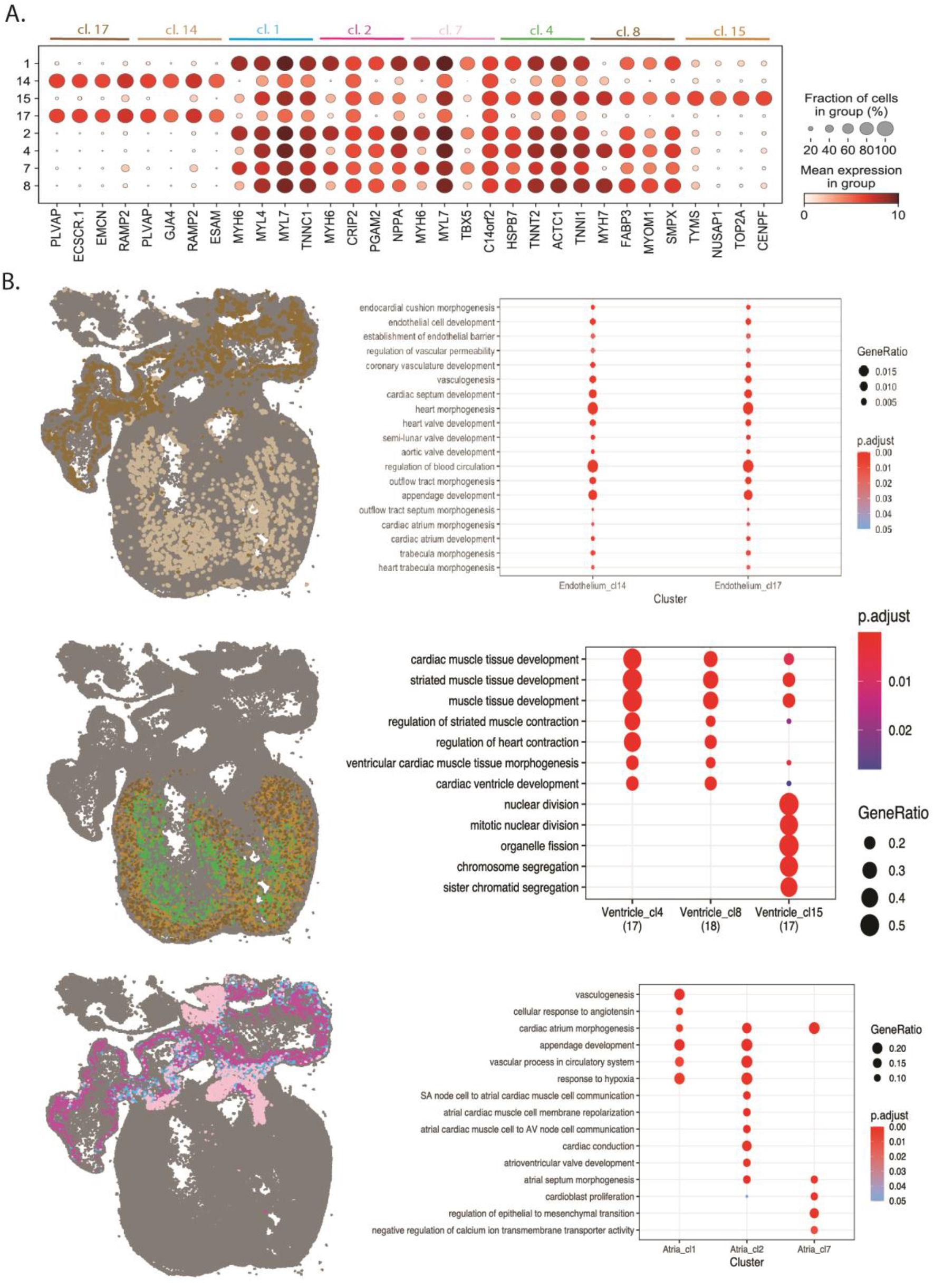
Exploration of new clusters identified within cardiomyocytes and endothelial cells. A. Dotplot representing the expression of the 4 most differentially expressed genes of each of the clusters related with endothelial cells (cluster 14 and 17), atrial (cluster 1,2 and 7) and ventricular (cluster 4, 8 and 15) cardiomyocytes. Expression is shown in the endothelial and cardiomyocyte related clusters linked to specific populations within the scRNA-seq dataset from Asp.et al.10 B. Spatial maps highlighting the reads assigned to the clusters related with endothelial cells and cardiomyocytes (right), with gene ontology enrichment of biological processes for top 15 most differentially expressed genes of each cluster (left). Color codes as in A.

High diversity was also found within cardiomyocytes, where three different populations were described within atrial cardiomyocytes and a total of five populations were found within ventricular cardiomyocytes, three of them having supporting scRNA-seq data **(Figure 2B)**. Moreover, spage2vec clusters present a better-defined region-specific location compared to analogous pciSeq cell-type maps in Asp et al., where some atrial cells are misplaced in the ventricles and vice versa (Supplementary Figure 4D).

Within ventricular cardiomyocytes, the five different spage2vec clusters defined presented unique expression patterns and spatial distributions from the periphery to the interior of the heart. However, not all the clusters were aligned with corresponding cell subpopulations from scRNA-seq data integration (Method). While clusters 4, 8 and 15 aligned within both ventricular and MYOZ2-enriched cardiomyocytes, cluster 16 and 24 presented a very weak alignment within the cell population sampled for scRNA-seq **(Figure 2B)**. Clusters aligning with specific cell subpopulations were further characterized **(Figure 3D, E)**. Cluster 4 had a location within the ventricular wall, was found to have a high expression of MYH7 and presented characteristics of trabecular myocardium while cluster 8 had an outer location and also had a strong expression of MYH7 consonant with outer, compact myocardium **(Figure 3A)**. Both cell types had GO characteristics of contracting ventricular muscle although these GO terms were more pronounced for trabecular myocardium. Cluster 15, which was smaller in size, expressed genes and GO characteristics of cell division and was preferentially located diffusively in the outer compact myocardium. These cells may thus be cardiomyoblasts participating in the consolidation of the compact myocardium.

Regarding atrial cardiomyocytes, the three clusters identified were found to have a different location **(Figure 3F)** and their specific markers were associated with distinct biological processes as suggested by Gene Ontology analysis **(Figure 3G)**. Cluster 1 was located mainly in the periphery of the atria and has GO characteristics of appendage formation while cluster 2 partly had a more central location and also expressed GO characteristics of cardiac conduction. Whereas cluster 7 was localized in the cranial and caudal part of the atria and had GO characteristics of morphogenesis and epithelial to mesenchymal transition. These spatial and GO characteristics are in keeping with the formation of the atrial septum that occurs at this stage of development.

Apart from the diversity found within cardiomyocytes, one of the most remarkable aspects of the analysis was the identification of different very thin sub-epicardial mesenchymal cell layers in the time point-specific analysis of pcw 6.5, possibly originating from epithelium via epithelial–mesenchymal transition (EMT)^15,16^. We described the expression signature of these clusters using diffusion maps and pseudotime analysis **(Figure 4A, B)**. The diffusion map suggests different differentiation processes involving the different mesenchymal clusters. By setting the root in epicardial cells (cluster 18), we identified two main branches in the pseudotime analysis, which could be indicating two differentiation paths involving epicardial cells: one involving the differentiation of epicardial cells into epicardial derived cells and fibroblasts (i.e., cluster 18-21-26-9-12-10) and a second one involving the possible differentiation of epicardial cells into atrial cardiomyocytes (i.e., cluster 18-11-1-2-7) **(Figure 4A, C)**. An additional branch connects epicardial derived cells to atrial cardiomyocytes (i.e., cluster 12-27-7-2-1), suggesting that EPDC undergo mesenchymal transition and differentiate into cardiomyocytes^17,18^. By mapping both the spage2vec identities and the pseudotime scores of these branches into the tissue we observed that the pseudotime described has a clear spatial component, matching with gradient from the periphery to the interior of the heart in the developing atria **(Figure 4D)**. GO analysis of clusters presenting enough supporting scRNA-seq cells show terms enriched for EMT and atrial morphogenesis **(Figure 4E)**.

**Figure 4.**
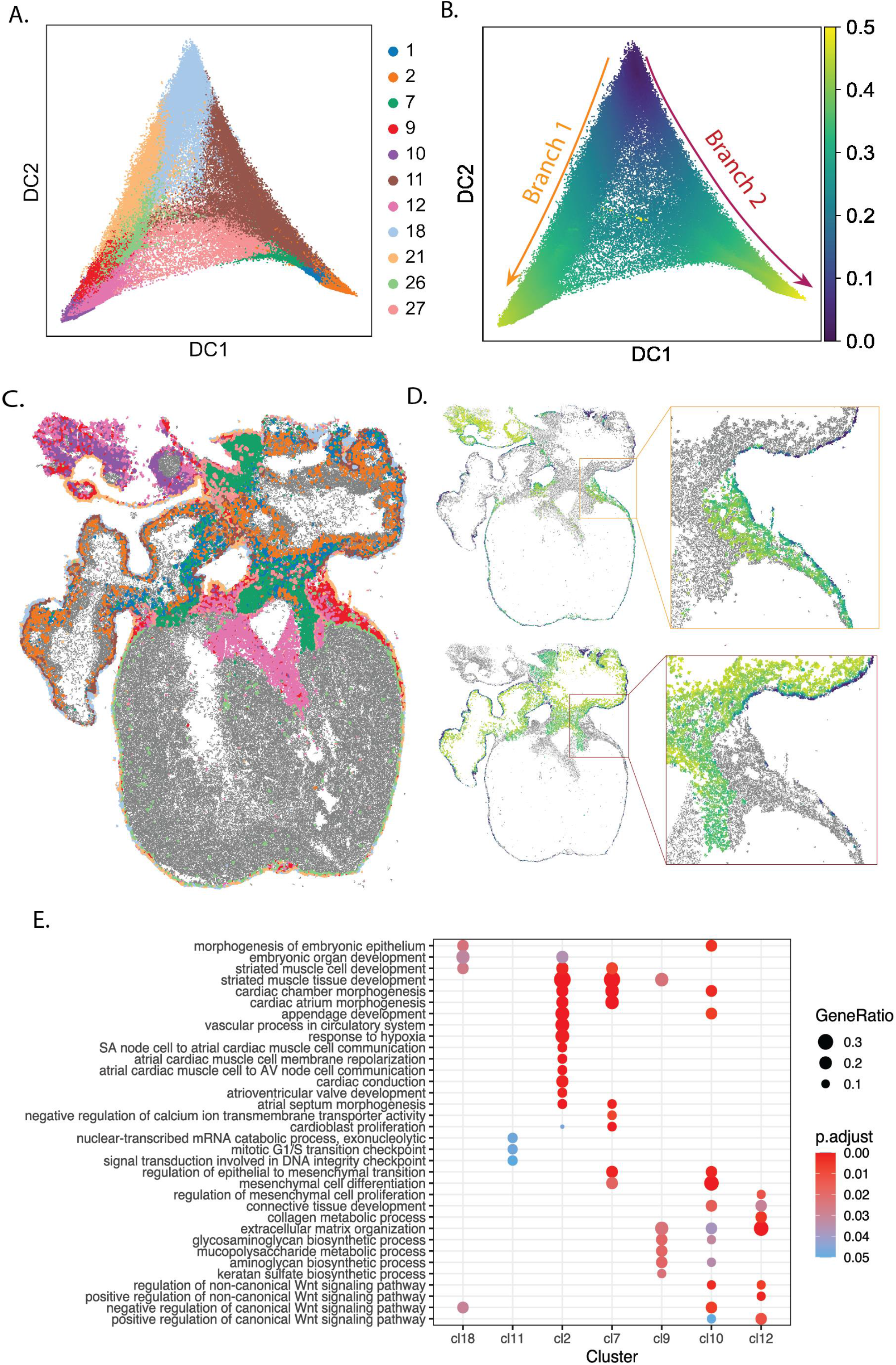
Description of the differentiation of epicardial cells in the human heart development. A,B. Diffusion map of pseudo-cell expression profiles defined in pcw 6.5 (Methods) and assigned to clusters related with epicardial cells, ventricular cardiomyocytes, epicardium-derived cells and fibroblasts. Each spot is labelled in A according to the cluster it was assigned to in Figure 2A. In B, the color of each spot represents its pseudotime score, considering the root in cluster 18 (epicardial cells). Two main branches can be observed. Pseudotime scores above 0.5 were trimmed for visualization purposes C. Spatial map highlighting the spots assigned to the clusters present in Figure 4A in one of the two sections from pcw 6.5. D. Spatial map representing the pseudotime scores of each of the spots described in Figure 4B in one of the sections from pcw 6.5 and a region of interest present in the same tissue. Clusters were represented in two different plots, depending on whether they were situated in Branch 1 (top) or Branch 2 (bottom) according to Figure 4A-B. E. Dot plot showing enrichment of Gene Ontology biological processes for top 15 most differentially expressed genes in the clusters represented in Figure 4A.

## DISCUSSION

The improvement of targeted spatially resolved transcriptomic approaches ^19,20^ in terms of signal-to-noise ratio, sequencing depth, number of genes and number of cells analyzed is leading towards the generation of larger datasets that will enable more and more comprehensive data driven spatial analysis. So far, methods such as ISS have primarily been a useful complement to scRNA-seq strategies by uncovering the spatial location of scRNA-seq defined cell populations. However, spatial molecular organization in itself presents intrinsic critical information of the cellular heterogeneity that is not captured by non-spatial methods, thus *de novo* approaches that do not rely on previous knowledge are starting to gain relevance in the field due to their notable advantages^13,21^. In this study, thanks to one of these *de novo* approaches, spage2vec^13^, we have been able to define 27 molecular signatures conserved during the developmental process of the heart based solely on the spatial location of the expressed molecules of 69 targeted genes.

In contrast with the original study^10^, where cell typing was constrained by availability of scRNA-seq data, our approach is able to define, in a spatiotemporal manner, different molecular signatures conserved through the different time points analyzed during the heart development. Our analysis showed to be especially relevant for capturing stable cell populations conserved through the developmental process, such as epicardial cells, and could be used for understanding biological processes like migration and differentiation. Supervised cell typing approaches^11,22,23^ will force the ISS data to fit signatures designed from scRNA-seq, with the risk of introducing biases and losing part of the potential biological information available in the ISS data. Furthermore, supervised approaches may fail to assign cells to a cell type due to discrepancies between the detected molecular signatures and the scRNA-seq data. As a consequence, while *de novo* approaches such as spage2vec assign a molecular signature to each read analyzed, probabilistic cell typing approaches avoid assigning a signature to many of the reads analyzed, missing in some cases molecular patterns with a true biological implication.

Moreover, unlike most existing cell typing strategies, spage2vec does not rely on cell segmentation. This aspect is highly beneficial when working with compact tissue, where cell borders are difficult to define. Spage2vec directly clusters the mRNA reads based on their local environment, and neighborhood information is incorporated in the process. Therefore, it is possible to discern populations that are similar in gene expression but have distinct spatial contexts in the tissue. In order to capture spatial signatures at cellular resolution, spage2vec aggregates local information from neighborhoods within a radius of 14.59 μm, which is a reasonable inter-cell distance, although the detected spatial clusters can represent cellular and even subcellular gene expression patterns. Since the method is completely unsupervised, super-cellular or sub-cellular patterns may also be captured depending on multiple factors that are related to the gene panel selected, sequencing resolution, and local differences in cell density.

For its unsupervised analysis, spage2vec depends on a targeted ISS gene panel. In this case, the genes were selected at an early stage of the Asp work^10^, based on scRNA-seq and Spatial Transcriptomics data. Despite the clear limitation of using a subset of markers for identifying clusters *de novo*, we have shown that leveraging deep learning representation power, spage2vec can also identify subpopulations through non-linear aggregation of spatial marker features, even without marker genes that can directly identify all cell populations. This is demonstrated by the identification of distinct atrial and ventricular subclusters with discrete GO characteristics. Another important cluster identified in spage2vec is the endocardial cluster that in Asp et al.^10^ was not subclustered out of the large scRNA-seq cluster comprising endothelial cells.

Apart from its ability to capture specific subpopulations, here we prove that spage2vec can be used to describe differentiation processes, including its spatial component. In this manuscript we report two main trajectories involving epicardial cells in atrial development. In fact, this observation is supported by Singh et al. 2013^24^, Greulich et al. 2011^25^, Cai et al. 2008^17^ and Zhou et al. 2008^18^, who report that at the atrial level epicardial cells flow into the atrial myocardial wall of venous origin and through epithelial-to-mesenchymal transition differentiate into arterial endothelium, smooth muscle and perivascular fibroblasts and may contribute to myocardialization of the atrial wall.

All in all, by applying spage2vec to study the human heart development we have been able to perform a spatiotemporal analysis of the cells found in post conception week 4.5-5, 6.5 and 9, identifying different molecular signatures within cardiomyocytes as well as an atrial subepicardial cell type previously unreported. This study shows the advantages of using *de novo* strategies that do not rely on cell segmentation and scRNA-seq to characterize developmental processes and opens the possibility of applying this approach to similar biological systems where reference single cell RNA sequencing data may be limited or not available.

## ACKNOWLEDGEMENTS

We thank Fredrik Nysjö and Christophe Avenel at the BioImage Informatics Facility, funded by SciLifeLab, the National Microscopy Infrastructure, NMI (VR-RFI 2019-00217), and the Chan-Zuckerberg Initiative, for help and support on data visualization. We’d also like to thank Christoffer Mattsson Langseth for his contribution in designing the study. The project was funded by the European Research Council via ERC Consolidator grant 682810 and Swedish Foundation for Strategic Research (grant BD150008), the Chan Zuckerberg Initiative, an advised fund of Silicon Valley Community Foundation; Erling-Persson Family Foundation; Knut and Alice Wallenberg Foundation; Swedish Research Council [2019-01238].

## AUTHOR CONTRIBUTIONS

X.Y. generated and applied the spage2vec models. S.M.S annotated the data with the help of C.S. and G.P.. G.P. performed the pseudotime analysis and the SpaGE integration. C.S. performed GO analysis and helped in interpreting the results. S.M.S. and G.P. coordinated the study. G.P., S.M.S., M.N. and C.W. conceived the study. S.M.S. designed and prepared figures. All authors contributed to writing and revising the manuscript.

## METHODS

### Data and code availability

All the code used to perform spage2vec on the developing human heart ISS data can be found in the following site: https://github.com/wahlby-lab/spage2vec_heart.

An online TissUUmaps^26^ viewer for interactive exploration of the analysis results can be found in: https://tissuumaps.research.it.uu.se/human_heart.html. All the data generated in this study can be downloaded from the TissUUmaps viewer for further exploration.

### Datasets

The ISS dataset of the developing human heart ^10^ comprises gene expression information of 69 marker genes and decoded spatial coordinates of mRNA spots in eight tissue sections at three developmental time points (Figure 1A). There are 189541, 812808, and 1471602 mRNA reads at the three time points respectively, summing up to a total of 2473951 reads.

### Spatiotemporal representation of ISS gene expression data with spage2vec

Spage2vec^13^ learns to map local neighborhood relationships between mRNA spots as distances in a continuous latent space using a deep learning model. As a result, a numerical vector is assigned to each individual mRNA spot describing its neighborhood composition. Therefore, molecules that share similar local environments are described with numerically similar vectors and consequently mapped in close proximity in the learnt latent space. In such a way, we are able to build a spatiotemporal representation of the spatial gene expression in an unsupervised manner and without using any prior information. The learnt representation is then used to perform clustering analysis in order to define localized gene expression signatures that represent cell-type signatures across the three embryonic stages.

### Constructing a spatial gene expression graph

We first construct a graph where each node represents an mRNA spot, with a one-hot encoding feature vector representing its corresponding gene. Each node is then connected by edges to its spatial local neighbors of the same tissue section within a maximum distance (d_max = 44.9 pixels/ 14.58 μm). We estimate the maximum distance such that 99% of nodes in the graph are connected to at least one neighbor. Connected components with less than six nodes are successively removed from the graph in order to exclude spurious reads such as spots located outside of the region of the heart sample, thus leaving 97.7% of the original mRNA reads for further processing.

### Graph neural network model and training

We train then a graph neural network on the spatial gene expression graph to produce the spage2vec latent representation for each mRNA spot. The neural network consists of two GraphSAGE^27^ layers. At each layer, the features of a node and its local neighborhood are aggregated and propagated to the next layer. The neural network learns its parameters in an unsupervised setting by minimizing a loss function based on random walks. The loss function of a node encourages similarity between the node and a direct neighbor that occurs in a random walk, and dissimilarity between the node and another node randomly sampled from the graph. Regarding the hyperparameters of the model, we use the mean aggregator at each layer and ReLU as activation function for the first layer. The size of each layer is 32. The model is trained for 10 epochs with a batch size equal to 64, using Adam optimizer^28^ with a learning rate equal to 0.001. The output for each mRNA spot is then a spage2vec latent vector of length 32.

### Cluster analysis and visualization

After predicting a latent vector for each mRNA spot based on its neighborhood composition, we compute a kNN (k = 15) weighted graph of the spage2vec latent vectors and apply the Leiden clustering algorithm^29^ (with clustering resolution r=1) on the kNN-graph. We then use PAGA^30^ to quantify the connectivity of acquired clusters, which represents the proximity of the clusters in the latent space. Each cluster with less than 1000 nodes is merged into the closer larger cluster in the PAGA graph having the maximum connectivity to the smaller cluster, if the connectivity was greater than 0.1. Otherwise, they are considered outliers and filtered out. After merging and filtering out the small clusters, we count the number of spots per cluster per gene followed by cluster-wise Z-score normalization to create a cluster expression matrix. This led to the final set of spage2vec clusters, which can be visualized interactively using TissUUmaps^26^.

### Spage2vec and scRNA-seq data integration

We perform data integration between spage2vec clusters of individual analysis of pcw 6.5 ISS data and the corresponding scRNA-seq data from Asp et al. Specifically, we first log-normalize scRNA-seq total counts per cell. Then, we generate pseudo-cell gene expression profiles for each mRNA spot by aggregating its k-nearest neighbor (k=100) in the spage2vec latent space. Next, we filter genes with less than 100 reads and log-normalize total counts per pseudo-cell. We thereafter integrate pseudo-cell and scRNA-seq gene expression profiles using SpaGE^14^. The two datasets are aligned by projecting them in a common latent space by domain adaptation^31^ using 30 principal vectors. After alignment, we can either infer the spatial profile of genes that are missing from the original ISS gene panel, or vice versa assign scRNA-seq cells to spage2vec clusters by k-nearest neighbor imputation.

Specifically, for each scRNA-seq cell we compute a cosine similarity in the common latent space with respect to all the k-th (k=15) nearest neighbor pseudo-cells, and we define correspondence with a spage2vec cluster as the sum of all cosine similarities with respect to those pseudo-cells belonging to the given spage2vec cluster. We then exclude scRNA-seq cells with low correspondence to spage2vec clusters (i.e. maximum cosine similarity smaller than 0.3), and we assign each scRNA-seq cell to the spage2vec cluster with highest cosine similarity. Spage2vec clusters with less than 10 scRNA-seq cells assigned are marked as weakly aligned as they miss enough supporting scRNA-seq cells and thus are excluded from further analyses.

## REFERENCES

1. The human body at cellular resolution: the NIH Human Biomolecular Atlas Program. Nature. Published online 2019. doi:10.1038/s41586-019-1629-x

2. Rozenblatt-Rosen O, Stubbington MJT, Regev A, Teichmann SA. The Human Cell Atlas: From vision to reality. Nature. Published online 2017. doi:10.1038/550451a

3. Macosko EZ, Basu A, Satija R, et al. Highly parallel genome-wide expression profiling of individual cells using nanoliter droplets. Cell. Published online 2015. doi:10.1016/j.cell.2015.05.002

4. Klein AM, Mazutis L, Akartuna I, et al. Droplet barcoding for single-cell transcriptomics applied to embryonic stem cells. Cell. Published online 2015. doi:10.1016/j.cell.2015.04.044

5. Moffitt JR, Hao J, Wang G, Chen KH, Babcock HP, Zhuang X. High-throughput single-cell gene-expression profiling with multiplexed error-robust fluorescence in situ hybridization. Proc Natl Acad Sci U S A. 2016;113(39):11046–11051. doi:10.1073/pnas.1612826113

6. Ke R, Mignardi M, Pacureanu A, et al. In situ sequencing for RNA analysis in preserved tissue and cells. Nat Methods. 2013;10(9):857–860. doi:10.1038/nmeth.2563

7. Codeluppi S, Borm LE, Zeisel A, et al. Spatial organization of the somatosensory cortex revealed by osmFISH. Nat Methods. 2018;15(11):932–935. doi:10.1038/s41592-018-0175-z

8. Eng CHL, Lawson M, Zhu Q, et al. Transcriptome scale super-resolved imaging in tissues by RNA seqFISH+. Nature. 2019;568(7751):235–239. doi:10.1038/s41586-019-1049-y

9. Ståhl PL, Salmén F, Vickovic S, et al. Visualization and analysis of gene expression in tissue sections by spatial transcriptomics. Science (80-). 2016;353(6294):78–82. doi:10.1126/science.aaf2403

10. Asp M, Giacomello S, Larsson L, et al. A Spatiotemporal Organ-Wide Gene Expression and Cell Atlas of the Developing Human Heart. Cell. 2019;179(7):1647–1660.e19. doi:10.1016/j.cell.2019.11.025

11. Qian X, Harris KD, Hauling T, et al. Probabilistic cell typing enables fine mapping of closely related cell types in situ. Nat Methods. 2020;17(1):101–106. doi:10.1038/s41592-019-0631-4

12. Phansalkar R, Red-Horse K. Techniques converge to map the developing human heart at single-cell level. Nature. Published online 2020. doi:10.1038/d41586-020-00151-z

13. Partel G, Wählby C. Spage2vec: Unsupervised representation of localized spatial gene expression signatures. FEBS J. Published online 2020. doi:10.1111/febs.15572

14. Abdelaal T, Mourragui S, Mahfouz A, Reinders MJT. SpaGE: Spatial Gene Enhancement using scRNA-seq. Nucleic Acids Res. 2020;48(18):E107-E107. doi:10.1093/nar/gkaa740

15. Kovacic JC, Mercader N, Torres M, Boehm M, Fuster V. Basic Science for Clinicians Epithelial-to-Mesenchymal and Endothelial-to-Mesenchymal Transition From Cardiovascular Development to Disease. Published online 2012. doi:10.1161/CIRCULATIONAHA.111.040352

16. Von Gise A, Pu WT. Review Endocardial and Epicardial Epithelial to Mesenchymal Transitions in Heart Development and Disease. Published online 2012. doi:10.1161/CIRCRESAHA.111.259960

17. Cai CL, Martin JC, Sun Y, et al. A myocardial lineage derives from Tbx18 epicardial cells. Nature. 2008;454(7200):104–108. doi:10.1038/nature06969

18. Zhou B, Ma Q, Rajagopal S, et al. LETTERS Epicardial progenitors contribute to the cardiomyocyte lineage in the developing heart. doi:10.1038/nature07060

19. Lee H, Salas SM, Gyllborg D, Nilsson M. Direct RNA targeted transcriptomic profiling in tissue using Hybridization-based RNA In Situ Sequencing (HybRISS). bioRxiv. Published online January 1, 2020:2020.12.02.408781. doi:10.1101/2020.12.02.408781

20. Gyllborg D, Nilsson M. HybISS: Hybridization based In Situ Sequencing. ProtocolsIo. Published online 2020. doi:10.17504/protocols.io.xy4fpyw

21. Park J, Choi W, Tiesmeyer S, et al. Segmentation free inference of cell types from in situ transcriptomics data. bioRxiv. Published online 2019. doi:10.1101/800748

22. Biancalani T, Scalia G, Buffoni L, et al. Deep learning and alignment of spatially-resolved whole transcriptomes of single cells in the mouse brain with Tangram. bioRxiv. Published online 2020.

23. Hie B, Bryson B, Berger B. Efficient integration of heterogeneous single-cell transcriptomes using Scanorama. Nat Biotechnol. 2019;37. doi:10.1038/s41587-019-0113-3

24. Singh MK, Epstein JA. Developmental Biology Epicardial Lineages and Cardiac Repair. J Dev Biol. 2013;1:1. doi:10.3390/jdb1020141

25. Greulich F, Rudat C, Kispert A. Mechanisms of T-box gene function in the developing heart. doi:10.1093/cvr/cvr112

26. Solorzano L, Partel G, Wählby C. TissUUmaps: Interactive visualization of large-scale spatial gene expression and tissue morphology data. Bioinformatics. 2020;36(15):4363–4365. doi:10.1093/bioinformatics/btaa541

27. Hamilton WL, Ying R, Leskovec J. GraphSAGE. Adv Neural Inf Process Syst. Published online 2017.

28. Kingma DP, Lei Ba J. ADAM: A METHOD FOR STOCHASTIC OPTIMIZATION.

29. Traag VA, Waltman L, van Eck NJ. From Louvain to Leiden: guaranteeing well-connected communities. Sci Rep. 2019;9(1). doi:10.1038/s41598-019-41695-z

30. Wolf FA, Hamey FK, Plass M, et al. PAGA: graph abstraction reconciles clustering with trajectory inference through a topology preserving map of single cells. Genome Biol. Published online 2019. doi:10.1186/s13059-019-1663-x

31. Mourragui S, Loog M, Van De Wiel MA, Reinders MJT, Wessels LFA. PRECISE: A domain adaptation approach to transfer predictors of drug response from pre-clinical models to tumors. In: Bioinformatics.; 2019. doi:10.1093/bioinformatics/btz372

