## Supplementary files 1-8 for "De novo spatiotemporal modelling of cell-type signatures identifies novel cell populations in the developmental human heart"

### SUPPLEMENTARY FIGURES 1-8

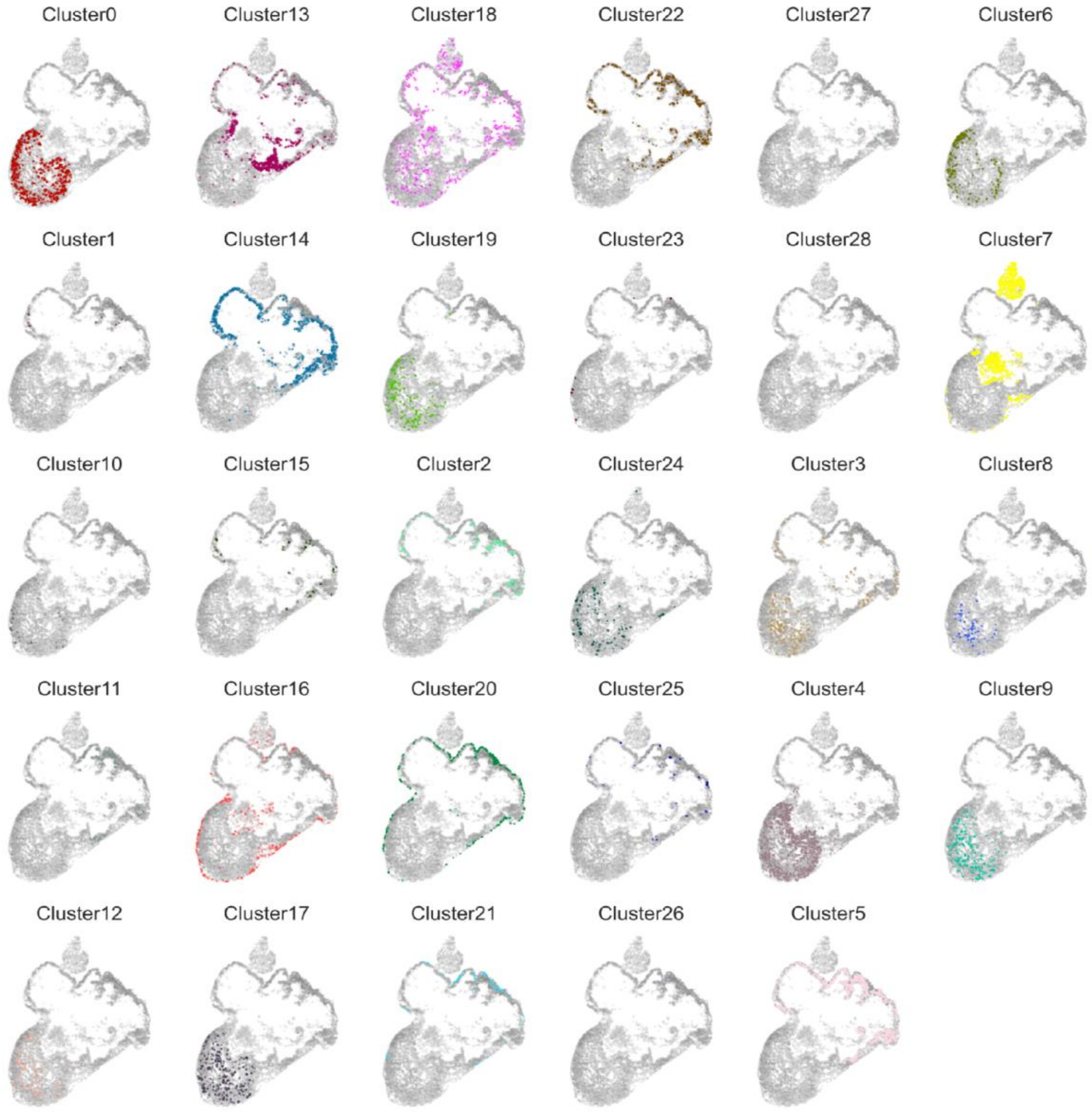


**Supplementary figure 1. Location of the spage2vec clusters in pcw 4.5-5.** Location of each of the spage2vec clusters described in Figure 1 in one of the three sections analyzed from pcw 4.5-5. Colors correspond to the ones used in Figure 1D. For interactive multi-resolution viewing: https://tissuumaps.research.it.uu.se/human_heart.html.

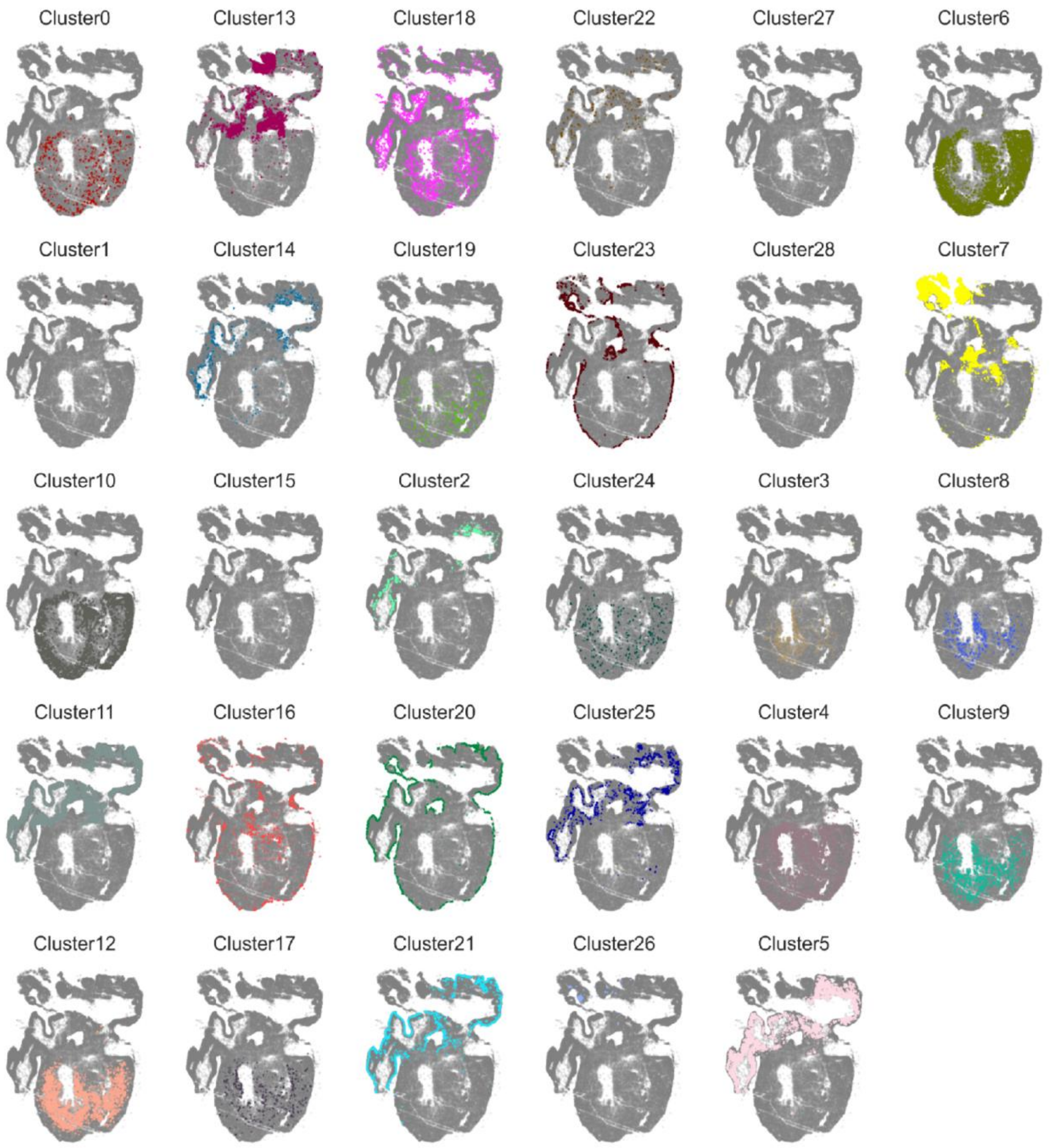


**Supplementary figure 2. Location of the spage2vec clusters in pcw 6.5.** Location of each of the spage2vec clusters described in Figure 1 in one of the two sections analyzed from pcw 6.5. Colors correspond to the ones used in Figure 1D. For interactive multi-resolution viewing: https://tissuumaps.research.it.uu.se/human_heart.html.

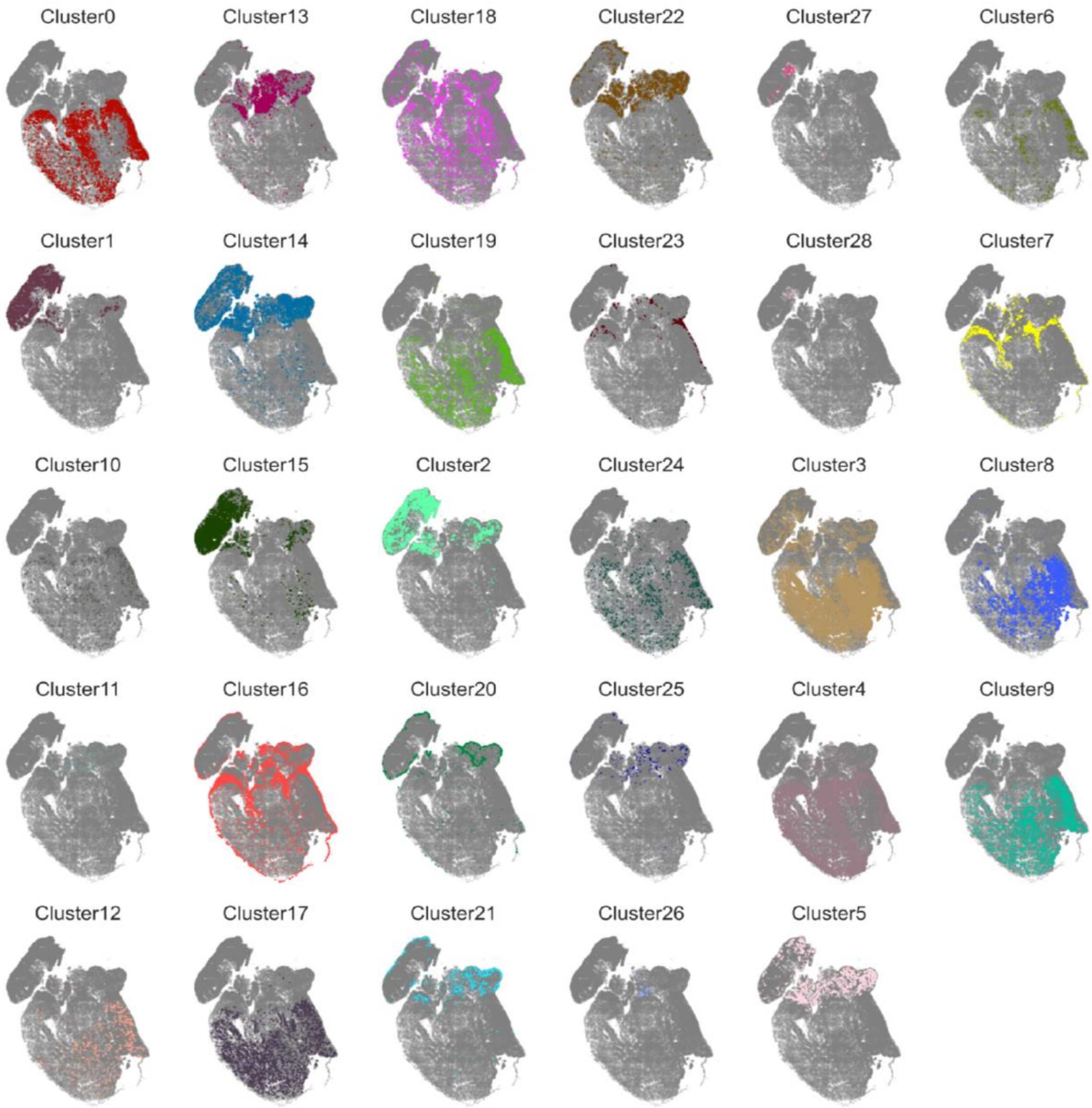


**Supplementary figure 3. Location of the spage2vec clusters in pcw 9.** Location of each of the spage2vec clusters described in Figure 1 in one of the three sections analyzed from pcw 9. Colors correspond to the ones used in Figure 1D. For interactive multi-resolution viewing: https://tissuumaps.research.it.uu.se/human_heart.html.

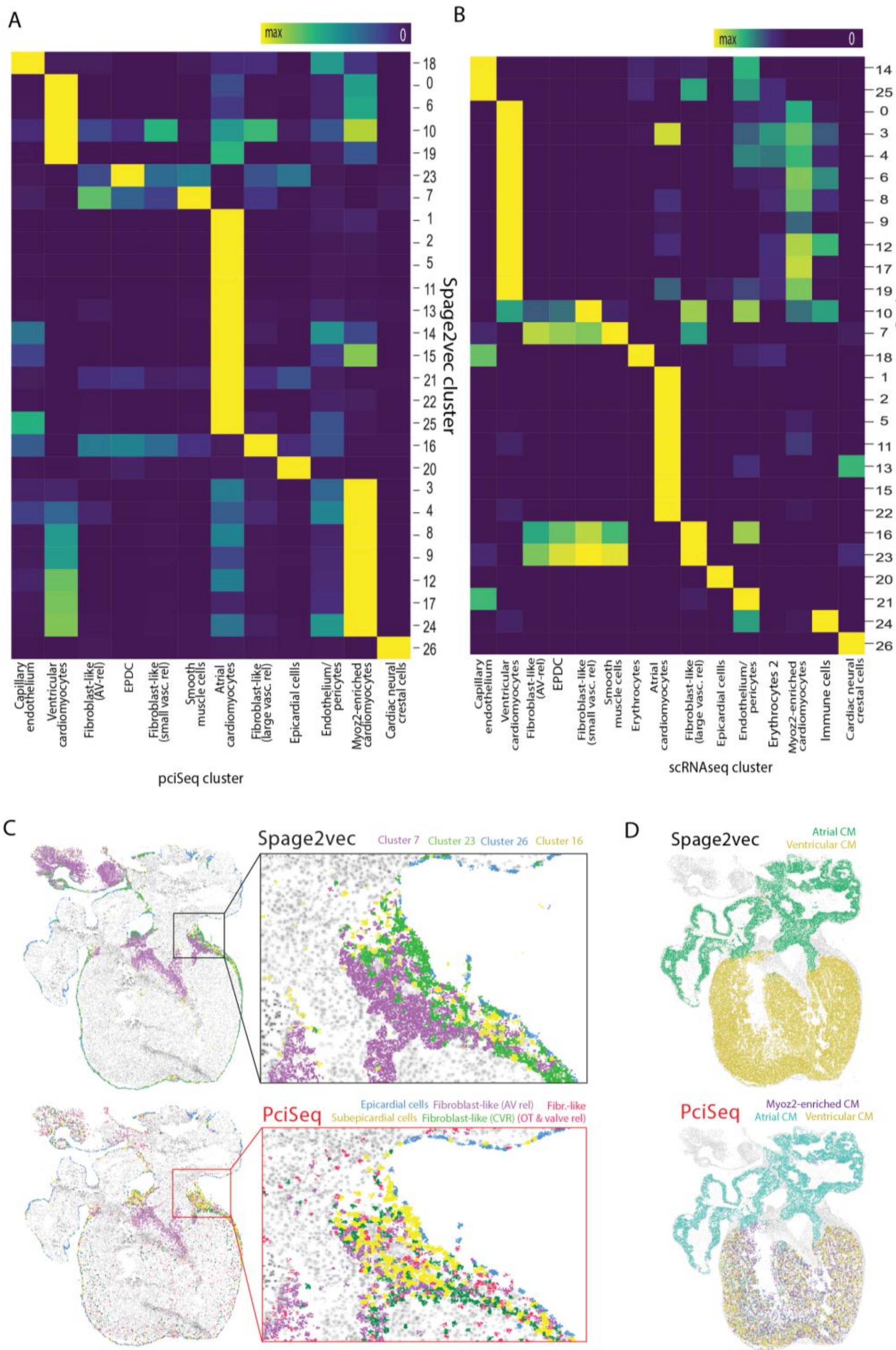


**Supplementary figure 4. Comparison of spage2vec clusters with cell-type annotations from Asp et al. A**. Heatmap representing the confusion matrix between the cell type assigned to each read via pciSeq in Asp et al.[10] and the spage2vec cluster annotations. **B**. Heatmap representing the correlation between the expression profile of each spage2vec cluster and each cell type described using scRNA-seq in Asp et al.[10] for the 69 genes included in both datasets. **C.** Spatial location of a subset of clusters from the spage2vec analysis (top) and pciSeq (bottom) in a specific sample from pcw 6.5. Clusters selected represent both epicardial cells and fibroblast-like cells /epicardium derived cells in both cases and colors have been based on the similarities between spage2vec clusters and pciSeq clusters. A zoomed in region is shown for both datasets. **D.** Spatial location of cardiomyocyte-related clusters defined by both spage2vec (top) and pciseq (bottom) in a specific section from pcw 6.5. Each spage2vec cluster assigned to cardiomyocytes were classified as atrial or ventricular according to their molecular signature in Figure 1C.

A

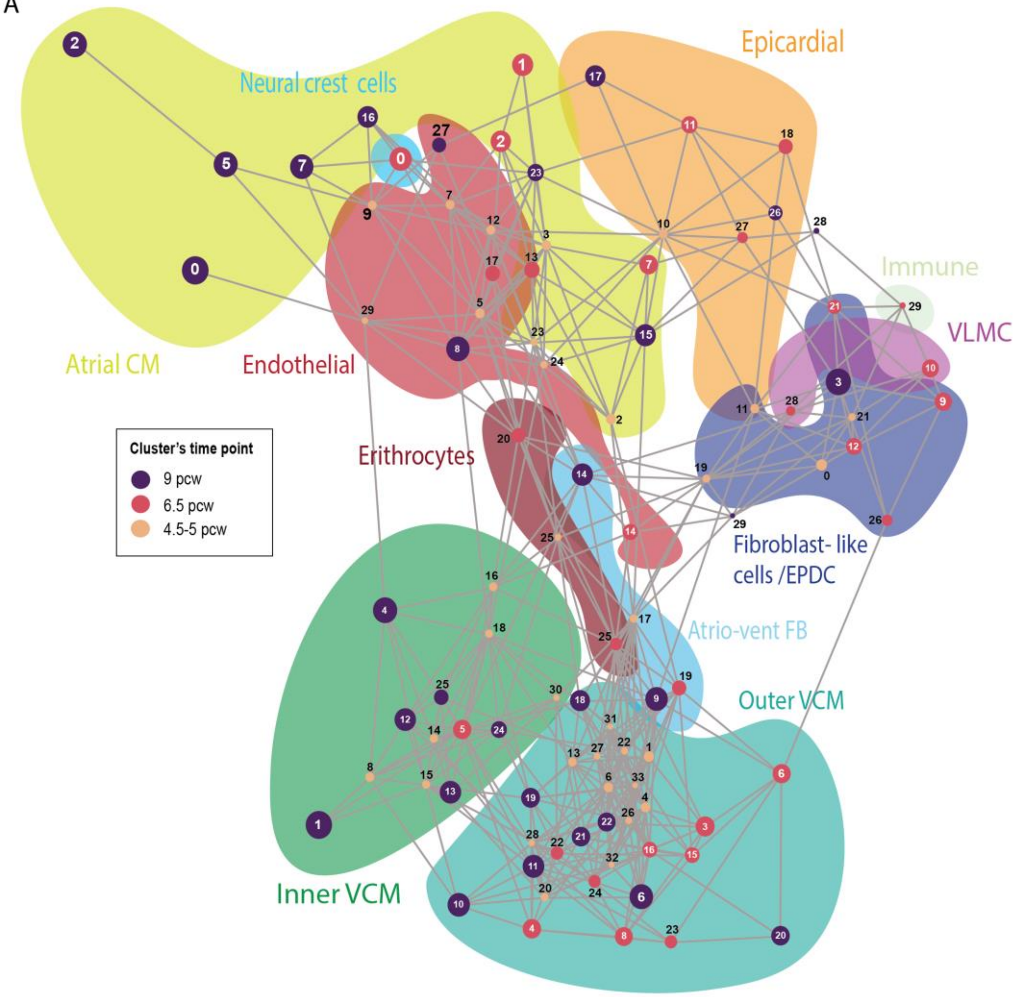


B

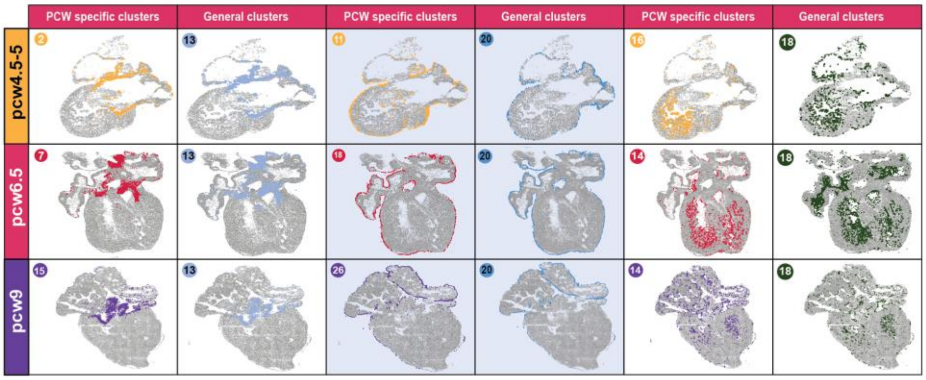

**Supplementary figure 5. Integration of time point-specific analyses. A.** PAGA plot representing all clusters found in the time-point specific analyses of pcw 4.5-5, pcw 6.5 and pcw 9. Each cluster is represented in a node and background colors indicate main cell type annotations. **B.** Spatial location's comparison between general clusters (Figure 1, Supplementary Figure 1-3) and time-point specific clusters. Three main clusters are represented: cluster 13 (left), cluster 20 (middle) and cluster 18 (right) in one of the samples of each time point, together with their correspondent time point-specific cluster.

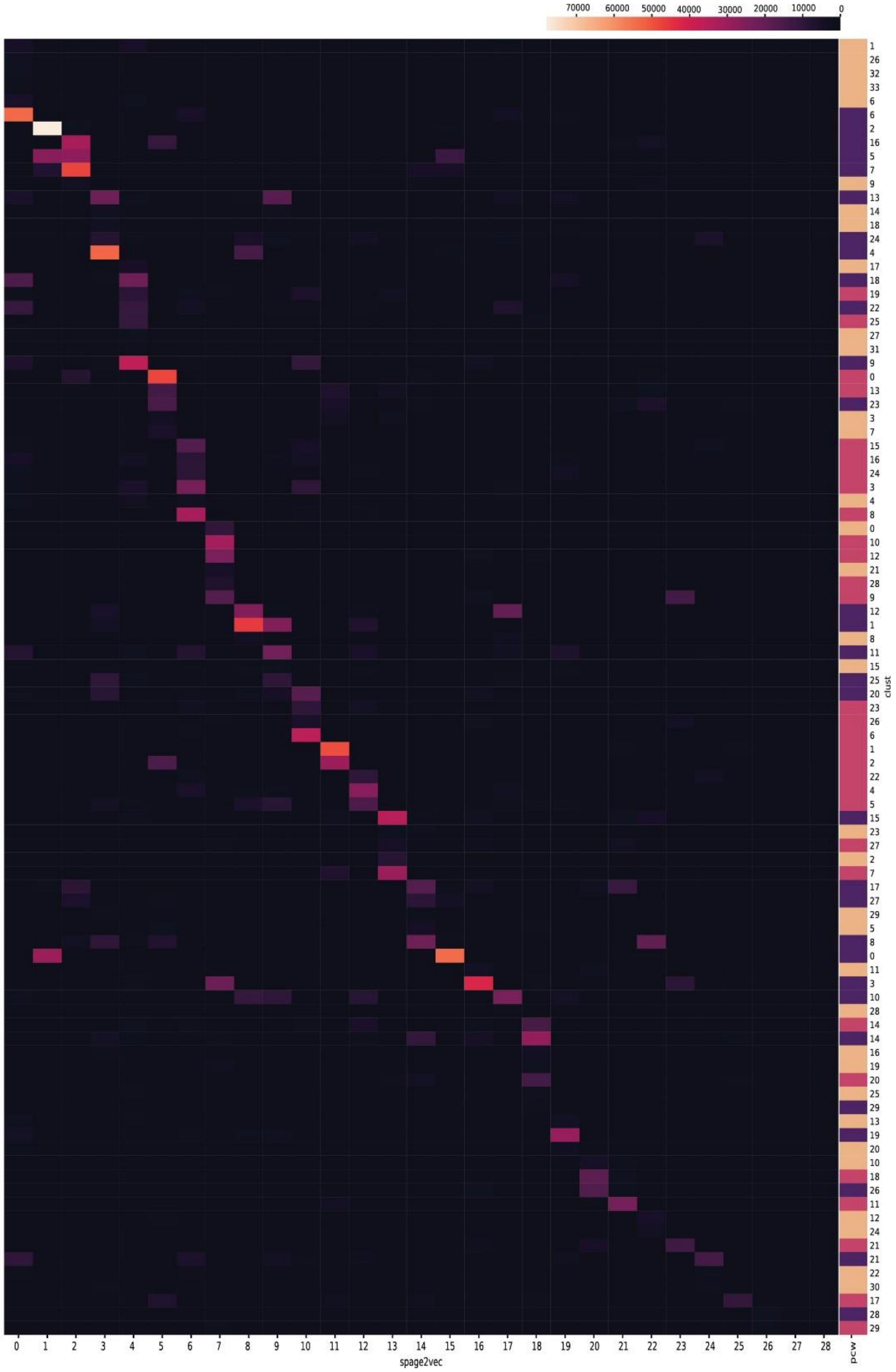


**Supplementary figure 6. Correspondence between general clusters and time point-specific cluster.** Heatmap representing the confusion matrix between the cluster assigned to each read in the general analysis (Figure 1, Supplementary Figure 1-3) and the cluster assigned to each read in the time point-specific analysis. Color column situated next to the time-point specific cluster labels indicates the time point where each cluster has been detected. The color code used in Figure 1A is used to label each time point.

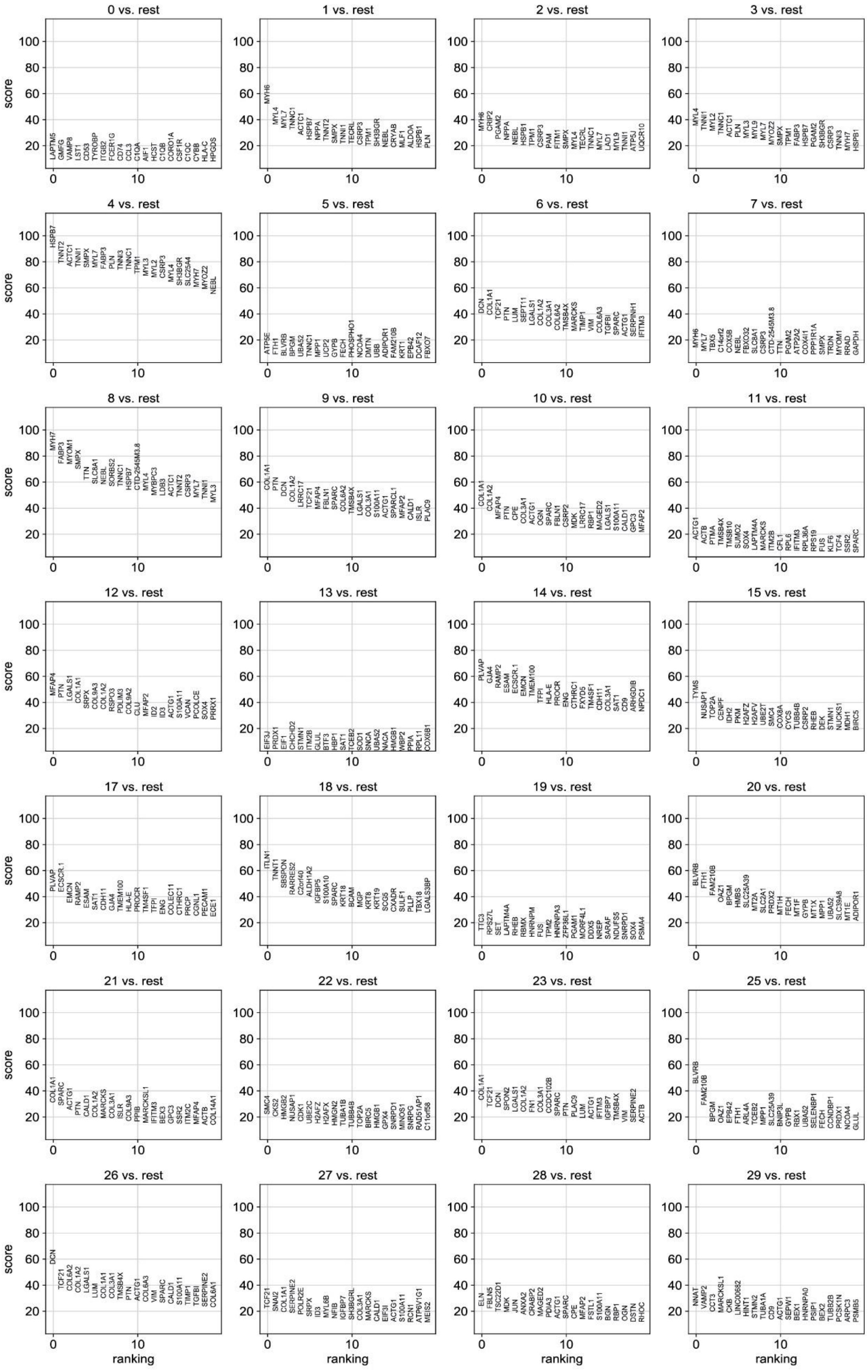
0 vs. rest
1 vs. rest
2 vs. rest
3 vs. rest
4 vs. rest
5 vs. rest
6 vs. rest
7 vs. rest
8 vs. rest
9 vs. rest
10 vs. rest
11 vs. rest
12 vs. rest
13 vs. rest
14 vs. rest
15 vs. rest
17 vs. rest
18 vs. rest
19 vs. rest
20 vs. rest
21 vs. rest
22 vs. rest
23 vs. rest
25 vs. rest
26 vs. rest
27 vs. rest
28 vs. rest
29 vs. rest
score
ranking

**Supplementary figure 7. Differential expression analysis of scRNA-seq data based on spage2vec cluster annotations.**Top 15 differentially expressed genes for the clusters found in the individual analysis of pcw 6.5. Scores of each gene (y axis) corresponds to the Wilcoxon rank-sum test score.

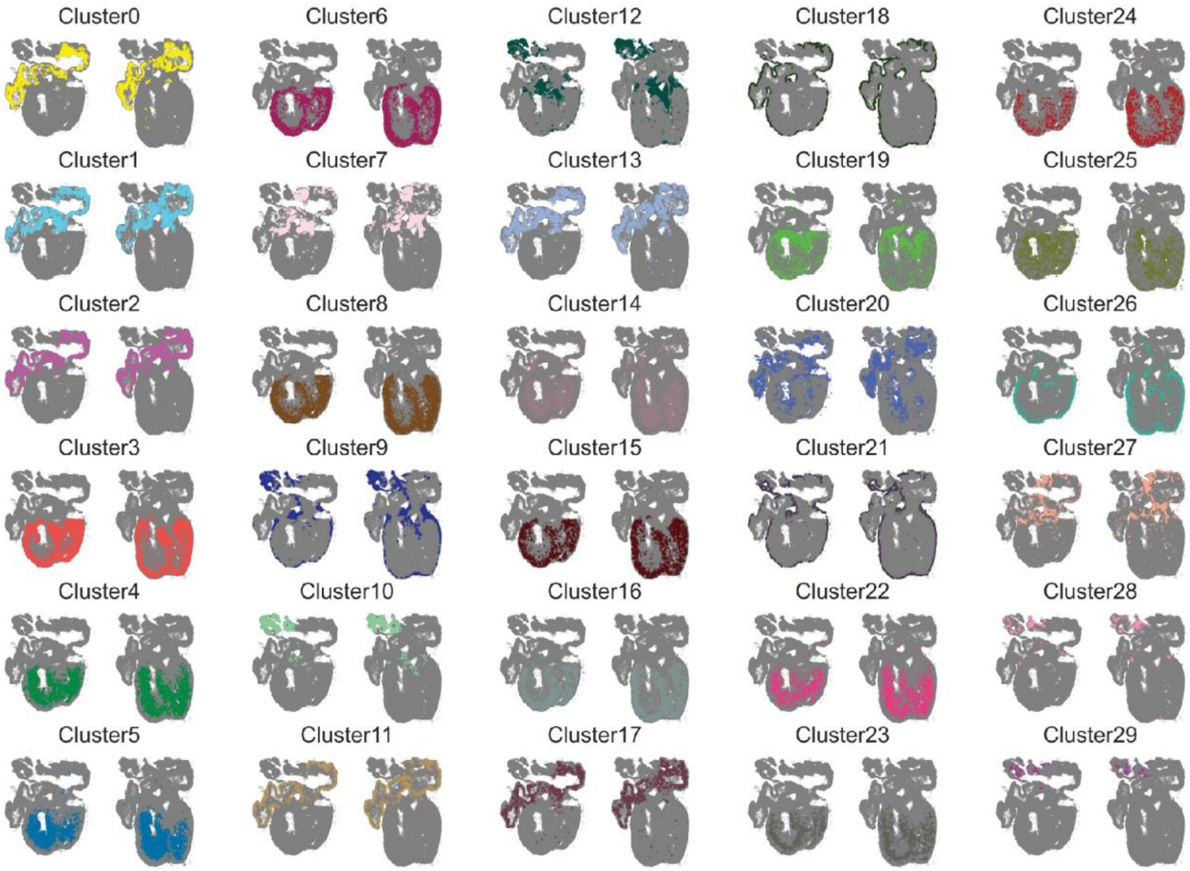


**Supplementary figure 8. Location of the spage2vec clusters in pcw 6.5 from the individual time point analysis.** Location of each of the spage2vec clusters described in Figure 2A in the two sections analyzed from pcw 6.5. Colors correspond to the ones used in the PAGA plots from Figure 2A. For interactive multi-resolution viewing: https://tissuumaps.research.it.uu.se/human_heart.html
